# Biophysical and enzymatic comparison of Bacillus safensis and Bacillus subtilis malate dehydrogenase (MDH) enzymes

**DOI:** 10.64898/2026.05.13.723581

**Authors:** Haley R. Zafiropoulo, Juliette E. Thomas, Nicholas R. Cortez, Korina Apostol, Alice de Sá, Ryan Khosravi, Liam Moore, Christopher E. Berndsen, Brianna Bibel

**Author notes:** These authors contributed equally (Korina Apostol, Alice de Sá, Ryan Khosravi, Liam Moore). Corresponding author contact information, Loyola Marymount University, Department of Chemistry and Biochemistry 1 LMU Dr. MS-8888, Los Angeles, CA 90045.

## Abstract

Species of *Bacillus* bacteria including *Bacillus safensis* and *Bacillus subtilis* are finding increasing uses in biotechnology and bioremediation, thanks in part to their metabolic robustness. Malate dehydrogenase (MDH) is at the heart of central metabolism and thus a better understanding of *Bacillus* MDH proteins could aid in the optimization of these applications. MDH of *Bacillus spp.* belong to the lactate dehydrogenase (LDH)-like class of MDH’s, otherwise known as the MDH3 class. Despite wide prevalence in nature among prokaryotes and archaea, this typically homotetrameric class is understudied compared to the MDH1 and MDH2 classes found in eukaryotes. We therefore recombinantly expressed and purified MDH proteins from two societally relevant *Bacillus spp.*–*B. safensis* and *B. subtilis*–and characterized them biophysically (via Size Exclusion Chromatography-Small Angle X-ray Scattering (SEC-SAXS) and Differential Scanning Fluorimetry (DSF)) and enzymatically (via spectroscopic activity assays). As expected based on their high sequence identity, the two MDH orthologs had similar properties in most regards, including a tetrameric structure and high susceptibility to substrate inhibition. However, we uncovered differences in conditional thermal stability, in addition to subtle differences in enzymatic activity that offer insight into the workings of LDH-like MDH.

**Summary statement:** Malate dehydrogenase (MDH) is a fundamental metabolic enzyme, from microbes to mammals, yet comparably little is known about microbial MDH, especially MDH of the tetrameric MDH3 class. We compare the biophysical and enzymatic properties of two such enzymes from the societally relevant bacterial species *Bacillus subtilis* and *Bacillus safensis*, offering useful insight with potential biotechnological implications.

## Introduction

Across all forms of life, metabolic enzymes enable cells to break down food for energy, build macromolecules, and safely process contaminants (among other functions). Although many core enzymes have been evolutionarily conserved, orthologs of these enzymes are often adapted to best meet the needs of the organisms they are in. One such fundamental enzyme is malate dehydrogenase (MDH), which facilitates the reversible interconversion of oxaloacetate and malate with the use of NAD(P)(H). MDH plays a crucial metabolic role in the citric acid cycle, fundamental to both catabolism (molecule-breaking) and anabolism (molecule-making). It also has additional roles in different organisms (e.g., facilitating the production of glucose via gluconeogenesis in eukaryotes). Dysregulation of human MDH has been linked to cancer (Cascón *et al*., 2015), encephalopathy (Ait-El-Mkadem *et al*., 2017), and other health conditions, making mammalian MDH the subject of increasing research. Bacterial MDH enzymes, on the other hand, have not been as extensively researched. They are, however, thought to have far more variation in molecular weight, enzymatic activity, and structural properties (Takahashi-Íñiguez *et al*., 2016a). Given the fundamental roles of bacteria in ecosystems and medicine (Singh, Shukla and Singh, 2024); their growing uses in biotechnology and bioremediation (Maglione *et al*., 2024; Morales-Mancera *et al*., 2025); and the rapid rise of protein engineering tools inspired by experimental data on extant proteins (Rennie and Oliver, 2025), characterization of bacterial MDH proteins has the potential to advance numerous fields.

Isoforms of MDH belong to an MDH/LDH (lactate dehydrogenase) superfamily whose members have evolved into distinct clades over millions of years (Wolyniak *et al*., 2024). MDH can be categorized into one of three different subgroups of MDH or LDH/MDH: MDH1, most commonly found in bacteria and eukaryotic cytoplasms; MDH2, primarily present in eukaryotic mitochondria; and MDH3 or LDH-like MDH, which is almost exclusively found in archaea and bacteria (Brochier-Armanet and Madern, 2021a). Within MDH lineages, structural adaptations have been identified that provide organism-specific properties while maintaining MDH activity. For example, the LDH-like MDH in the halophilic archaeon *Haloarcula marismortui* organizes surface amino acid residues within the solvation shell to allow high protein solubility and provide stabilizing interactions in high-salt environments (Richard *et al*., 2000). Meanwhile, thermophilic marine molluscs have evolved to have MDH1-class MDH proteins with loop regions that are less flexible than orthologous MDH proteins of cold-adapted molluscs (Dong *et al*., 2018).

Despite its broad distribution, comparatively little is known about MDH3 isoforms, which are thought to be primarily tetrameric, compared to the typically dimeric MDH1 & MDH2. The tetrameric structure may have implications for substrate/enzyme interactions, allosteric regulation, and protein stability (Kalimeri *et al*., 2014). Therefore, we set out to characterize and compare the biophysical and enzymatic properties of MDH3 isozymes from two societally relevant species of *Bacillus* bacteria: the model organism and biotechnology workhorse *Bacillus subtilis* (*B. subtilis* or *B. sub)*, which can be used for biomanufacturing (including of organic acids) (Liu *et al*., 2017), and *Bacillus safensis* (*B. safensis* or *B. saf*), which has undergone little research but has been proposed for use in heavy metal bioremediation due to its ability to thrive in harsh environments and reduce toxic hexavalent chromium, Cr(VI), to the benign Cr(III) (Harboul *et al*., 2023). We therefore performed spectroscopic enzyme activity assays to measure the kinetic parameters of *B. safensis* MDH (BsafMDH) and *B. subtilis* MDH (BsubMDH), differential scanning fluorimetry (DSF) to investigate their thermal stability, and size exclusion chromatography-small angle X-ray modeling (SEC-SAXS) to determine their oligomeric status and overall shapes. Our results suggest high similarity between BsafMDH and BsubMDH, as expected, but also reveal subtle enzymatic and structural differences that lay the groundwork for future investigation.

Given the requirement for metabolic robustness to carry out these processes and the central role of MDH in metabolism, a deeper understanding of MDH in these species could lend insight into metabolic adaptations that enable these processes and potentially allow for more efficient, environmentally friendly biotechnology and bioremediation applications. Additionally, given the high degree of similarity between the two MDH isozymes (∼94% amino acid identity) (Fig. 1), differences in their characteristics could offer direct insight into structural factors that affect LDH-like MDH activity.

**Fig 1.**
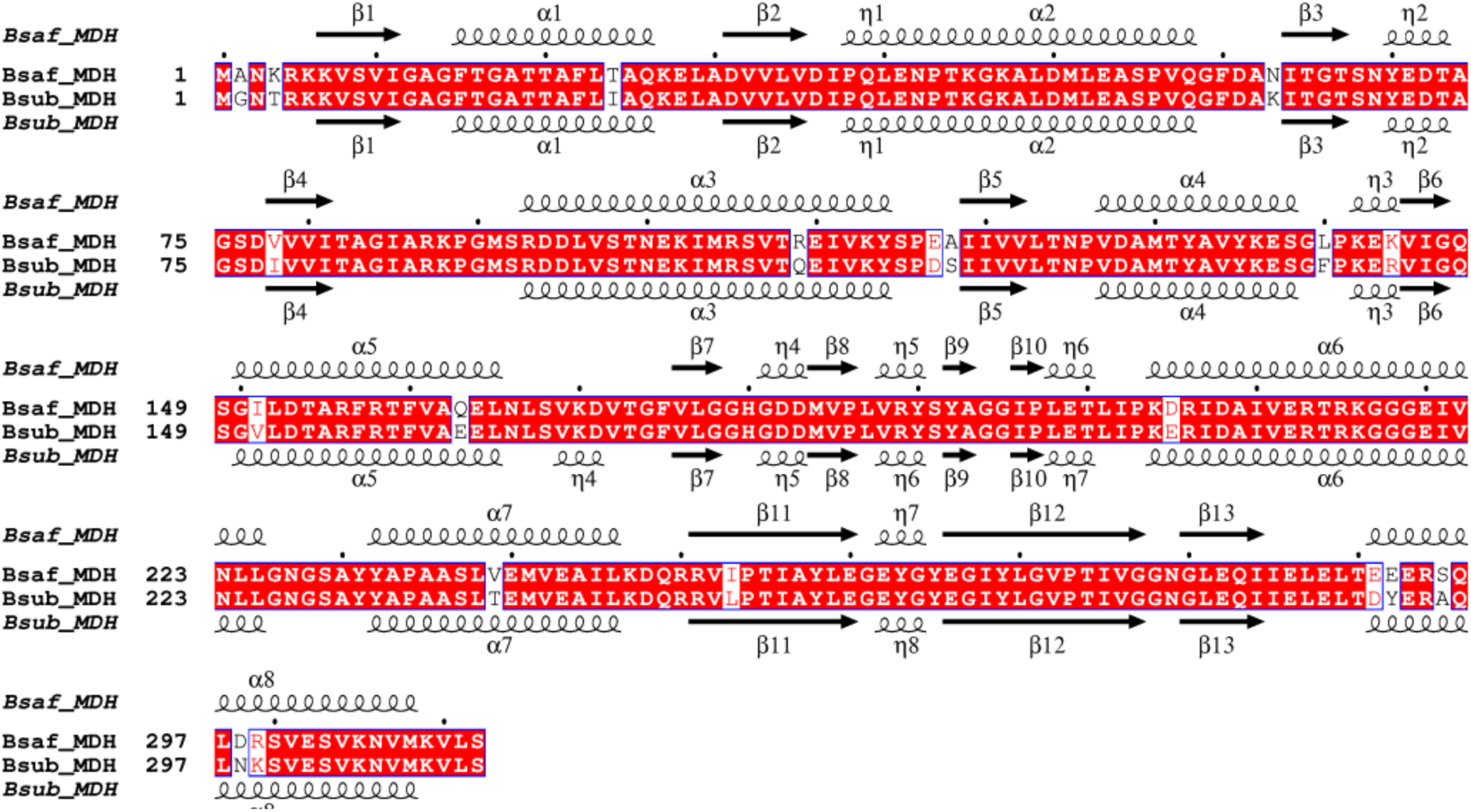
*Bacillus safensis* and *Bacillus subtilis* malate dehydrogenase (MDH) proteins have high (∼94%) amino acid sequence identity and predicted secondary structures. Alignment figure, colored by “% Equivalent,” was generated using ESPript (Gouet, Robert and Courcelle, 2003), incorporating secondary structure from AlphaFold 3 (Abramson *et al*., 2024) predictions based off of UniProt (The UniProt Consortium, 2025) accessions A0A0M2EAA6 (Bsaf_MDH) and P49814 (Bsub_MDH).

## Results

### *B. safensis* and *B. subtilis* MDH have highly similar tetrameric structures

We chose these two isozymes to study in part because the high sequence identity between them will allow for dissection of the biophysical roots of any differences in structure, enzymatic activity, and/or dynamics. As a first step in this direction, we set out to characterize and compare their biophysical properties. *Bacillus* MDH have generally been predicted to be tetrameric, based on phylogenetic membership in the MDH3 class, homology modeling, and, for some orthologs, experimental data (Murphey *et al*., 1967). To confirm the tetrameric structure of *B. safensis* and *B. subtilis* MDH, we expressed and purified recombinant *B. safensis* and *B. subtilis* MDH (Fig. S1) and performed SEC-SAXS analysis at pH 8.0 (Fig. 2 and Table 1).

**Fig. 2.**
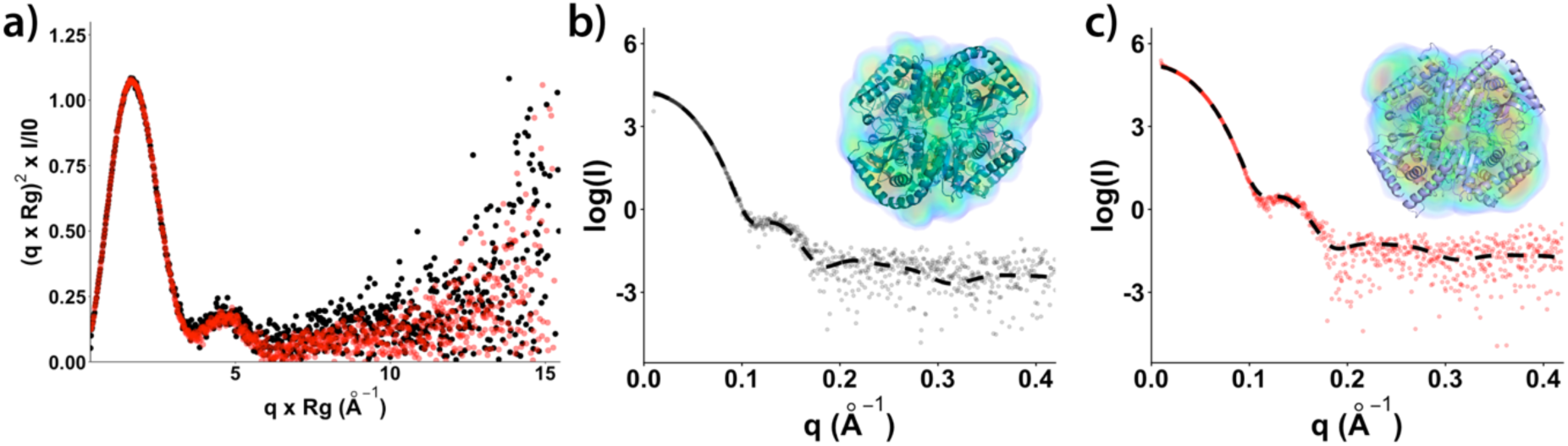
Small-angle X-ray scattering of BsubMDH and BsafMDH. **a)** Kratky plots comparing the shape of BsubMDH (black) and BsafMDH (red). **b)** FOXS fit of the BsubMDH SAXS data to the AlphaFold3 model (χ^2^ = 1.1) and fit of the model to the DENSS envelope (inset). **c)** FOXS fit of the BsafMDH SAXS data to the AlphaFold3 model (χ^2^ = 0.9) and fit of the model to the DENSS envelope (inset). The DENSS calculated real space correlation value for BsubMDH was 0.84 and for BsafMDH it was 0.79.

**Table 1:** SEC-SAXS data.

|  | **BsubMDH** | **BsafMDH** |
| --- | --- | --- |
| **Wavelength (Å)** | 1.127 | 1.127 |
| **q range (Å<sup>-1</sup>)** | 0.0098-0.4707 | 0.0099-0.4707 |
| **Concentration (mg mL<sup>-1</sup>)** | 5 | 5 |
| **Temperature (K)** | 283 | 283 |
| **I (0) (A.U.) (from p[r])** | 182 ± 0.2 | 69.5 ± 0.1 |
| **Rg (Å) from p[r])** | 32.3 ± 0.1 | 32.6 ± 0.1 |
| **I(0) (A.U.) (from Guinier)** | 181 ± 0.4 | 69.4 ± 0.2 |
| **Rg (Å) (from Guinier)** | 32.7 ± 0.1 | 32.9 ± 0.1 |
| **Dmax (Å)** | 97 | 93 |
| **Porod volume estimate**<br>**(Å<sup>3</sup>) (Vp)** | 1.8 x 10<sup>5</sup> | 1.76 x 10<sup>5</sup> |
| **Molecular mass Bayes**<br>**(kDa)** | 138.2 [93.8% C.I. 134.3 to<br>151.4] | 130.9 [97.4% C.I. 121.5 to 142.2] |
| **Molecular mass Mr (from**<br>**Porod volume) (kDa)** | 149.4 | 146.1 |
| **Calculated monomeric Mr**<br>**from sequence (kDa)** | 38.8 | 38.8 |
| **SASBDB ID** |  |  |
| **$\chi^2$ of FOXS fit model** | 1.1 | 0.9 |
| **Primary data reduction** | RAW | RAW |
| **Data processing** | RAW | RAW |
| **Computation of model intensities** | FOXS | FOXS |
| **Three-dimensional graphic representations** | ChimeraX and PyMOL | ChimeraX and PyMOL |

The experimentally determined molecular weights (MWs) obtained through SEC-SAXS of BsafMDH and BsubMDH MDH were 130.9 kDa (97.4% confidence interval (C.I.) of 121.5-142.2 kDa) and 138.2 (93.8% C.I. of 134.3-151.4 kDa), respectively (Table 1). These values are consistent with tetrameric structures as each monomer is calculated to have a MW of ∼35 kDa based on the sequence for a tetrameric weight of 140 kDa. Furthermore, Kratky plots suggested that, at least under these conditions, the overall shapes of the two isozymes were highly similar (Fig. 2a). FOXS fitting of AlphaFold3 models of the tetramer to the SAXS data resulted in fits with χ^2^ values of 1.1 for BsubMDH and 0.9 for BsafMDH (Figs. 2b, 2c, and Table 1). We further calculated the envelopes from the SAXS data using DENSS and aligned the data to the 3-D structures (Figs. 2b, 2c, & Table 1). The real space correlation for BsubMDH was 0.84 and for BsafMDH it was 0.79; both numbers suggest a reasonable match of the envelopes to the respective SAXS data. In sum, both computational modeling and biophysical data support *B. safensis* and *B. subtilis* MDH proteins having a nearly identical globular, tetrameric shape.

### *B. safensis* and *B. subtilis* MDH have similar enzymatic activity

We next set out to characterize the two proteins enzymatically, using conventional spectroscopic activity assays in which the disappearance (or appearance) of NADH is monitored via 340 nm wavelength absorbance. The MDH isoforms tested displayed similar kinetic properties in both the reductive (OAA + NAD^+^ + H^+^ → malate) and oxidative (malate + NAD^+^ → OAA + NADH + H^+^) directions (Fig. 3 and Table 2). *k_cat_* values were slightly higher for BsafMDH than BsubMDH with both NADH (153 ± 8 s^-1^ vs. 117 ± 4 s^-1^) and OAA (120 ± 10 s^-1^ vs 92 ± 6 s^-1^) as substrates, but not malate (21.6 ± 0.6 s^-1^ vs 21.6 ± 0.8 s^-1^). Compared to BsafMDH, BsubMDH had a slightly lower K_M_ for OAA, but not NADH or malate: K_M_ towards OAA (at pH 8.0) of 14 ± 3 µM for BsafMDH and 8 ± 2 µM for BsubMDH; K_M_ towards NADH of 10 ± 2 µM for BsafMDH and 14 ± 2 µM for BsubMDH; K_M_ towards malate of 61 ± 7 µM for BsafMDH and 69 ± 10 µM for BsafMDH. As is typical for MDH proteins in vitro (de Lorenzo *et al*., 2024b), the oxidative direction (malate to oxaloacetate, Fig. 3d) showed lower *k_cat_* values than the reductive direction (oxaloacetate to malate, Fig. 3a and 3b).

**Fig. 3.**
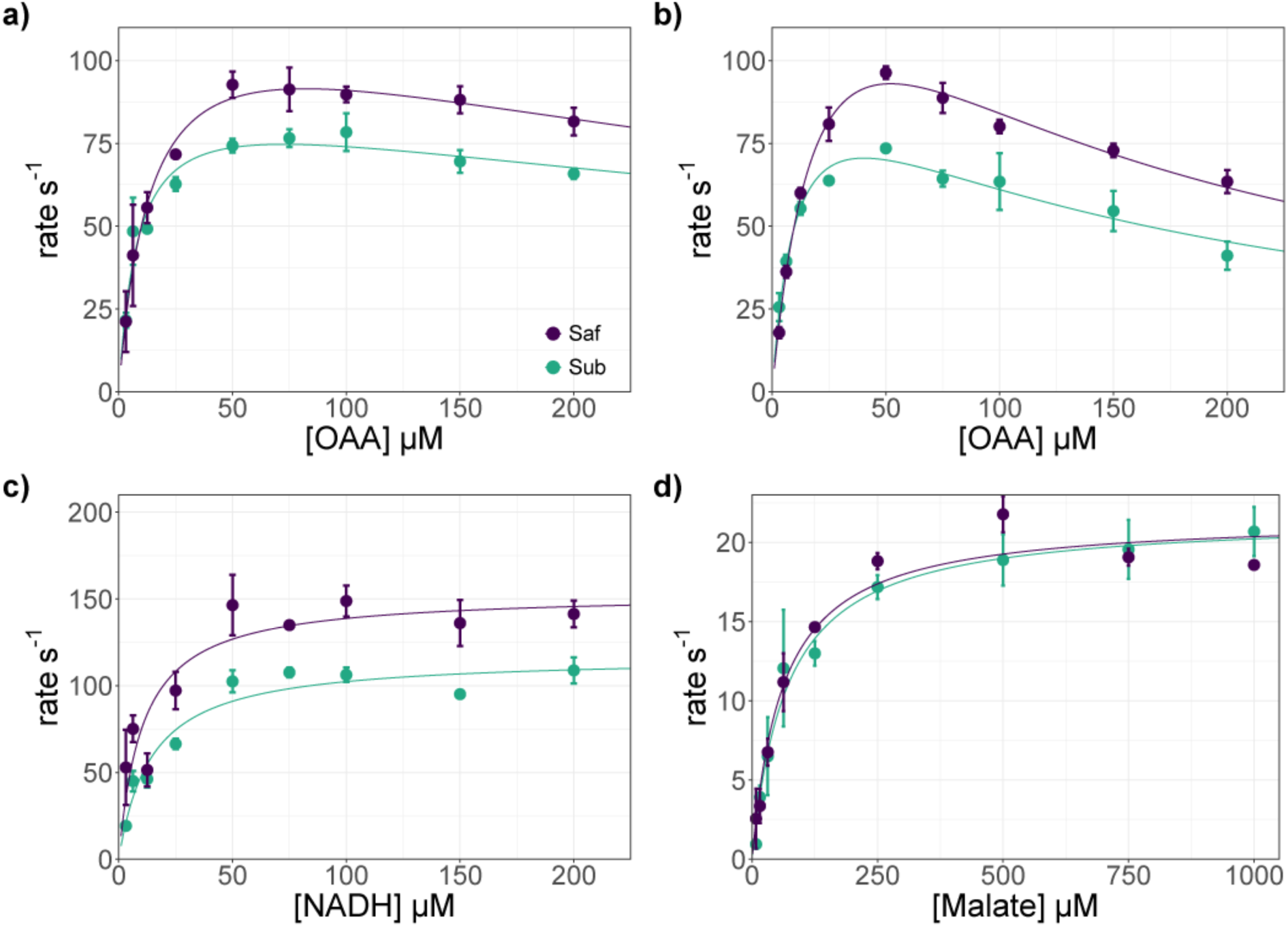
*B. safensis* MDH and *B. subtilis* MDH have similar kinetic properties. a-b) Kinetic assays performed in the reductive (oxaloacetate to malate) direction with a range of oxaloacetate (OAA) concentrations at pH 8.0 **(a)** and 7.4 **(b)** show similar kinetic properties, with pronounced substrate inhibition, especially at the lower pH. BsubMDH had a slightly lower K_M_ and *k_cat_* (See Table 2). Assays were run in 50 mM sodium phosphate buffer with NADH concentrations held at 100 µM. **c)** Kinetic assays performed in the reductive direction with a range of NADH concentrations at pH 8.0 show similar kinetic parameters between the isoforms, with nearly identical K_M_ values, but BsubMDH had a slightly lower *k_cat_*. Assays were run in 50 mM sodium phosphate buffer, pH 8.0 with OAA concentrations held at 100 µM. **d)** Kinetic assays performed in the oxidative direction with a range of malate concentrations at pH 8.0 show similar kinetic parameters between the isoforms. Assays were run in 50 mM sodium phosphate buffer, pH 8.0 with NAD^+^ concentrations held at 5 mM. Data is presented mean ± standard deviation, n = 3 technical replicates.

**Table 2:** Kinetic parameters of MDH enzyme activity.

| Oxaloacetate (OAA) |  |  |  |  |
| --- | --- | --- | --- | --- |
| pH 7.4 |  |  |  |  |
| | $k_{cat}$ (s<sup>-1</sup>) | $K_M$ (μM) | $k_{cat}/K_M$ (M<sup>-1</sup>s<sup>-1</sup>) | $K_i$ (μM) |
| **BsafMDH** | 180 ± 10 | 25 ± 3 | (7 ± 1) × 10<sup>6</sup> | 110 ± 10 |
| **BsubMDH** | 109 ± 9 | 11 ± 2 | (1 ± 0.2) × 10<sup>7</sup> | 150 ± 30 |
| pH 8.0 |  |  |  |  |
| | $k_{cat}$ (s<sup>-1</sup>) | $K_M$ (μM) | $k_{cat}/K_M$ (M<sup>-1</sup>s<sup>-1</sup>) | $K_i$ (μM) |
| **BsafMDH** | 120 ± 10 | 14 ± 3 | (9 ± 2) × 10<sup>6</sup> | 500 ± 200 |
| **BsubMDH** | 92 ± 6 | 8 ± 2 | (1.2 ± 0.3) × 10<sup>7</sup> | 600 ± 200 |
| NADH |  |  |  |  |
| | $k_{cat}$ (s<sup>-1</sup>) | $K_M$ (μM) | $k_{cat}/K_M$ (M<sup>-1</sup>s<sup>-1</sup>) | |
| **BsafMDH** | 153 ± 8 | 10 ± 2 | (1.5 ± 0.3) × 10<sup>7</sup> |  |
| **BsubMDH** | 117 ± 4 | 14 ± 2 | (8 ± 1) × 10<sup>6</sup> |  |
| Malate |  |  |  |  |
| | $k_{cat}$ (s<sup>-1</sup>) | $K_M$ (μM) | $k_{cat}/K_M$ (M<sup>-1</sup>s<sup>-1</sup>) | |
| **BsafMDH** | 21.6 ± 0.6 | 61 ± 7 | (3.5 ± 0.4) × 10<sup>5</sup> |  |
| **BsubMDH** | 21.6 ± 0.8 | 69 ± 10 | (3.1 ± 0.5) × 10<sup>5</sup> |  |

Consistent with reported findings from a variety of MDH homologs (Takahashi-Íñiguez *et al*., 2016b; Martinez-Vaz *et al*., 2024), we observed substrate inhibition by oxaloacetate, with K_i_ values of 500 ± 200 µM for BsafMDH and 600 ± 200 µM for BsubMDH (Fig 3a and Table 2). Substrate inhibition by OAA has been reported to be more severe at lower pH, theorized to be due to increased keto-enol tautomerization (Bernstein *et al*., 1978). Therefore, we additionally tested for OAA inhibition at pH 7.4. In this more acidic environment, BsafMDH and BsubMDH showed evidence of OAA inhibition at as low as 75 µM of OAA, with K_i_ values of 110 ± 10 µM for BsafMDH and 150 ± 30 µM for BsubMDH (Figure 3b and Table 2). Notably, 100 µM OAA concentration is frequently used for other isoforms in routine assays, with many reported values from assays performed at higher concentrations. Both the K_M_ and the *k_cat_* were higher at pH 7.4 compared to pH 8.0 for both BsafMDH (25 ± 3 µM and 180 ± 10 s^-1^ at pH 7.4 vs 14 ± 3 µM and 120 ± 10 s^-1^ at pH 8.0) and BsubMDH (11 ± 2 µM and 109 ± 9 s^-1^ at pH 7.4 vs 8 ± 2 µM and 92 ± 6 s^-1^ at pH 8.0). These differences were greater for BsafMDH than BsubMDH, but largely canceled out to give similar *k_cat_/*K_M_ values for the two enzymes ((7 ± 1) x 10^6^ M^-1^ s^-1^ for BsafMDH and (1.0 ± 0.2) x 10^7^ M^-1^ s^-1^ for BsubMDH) (Table 3).

**Table 3:**
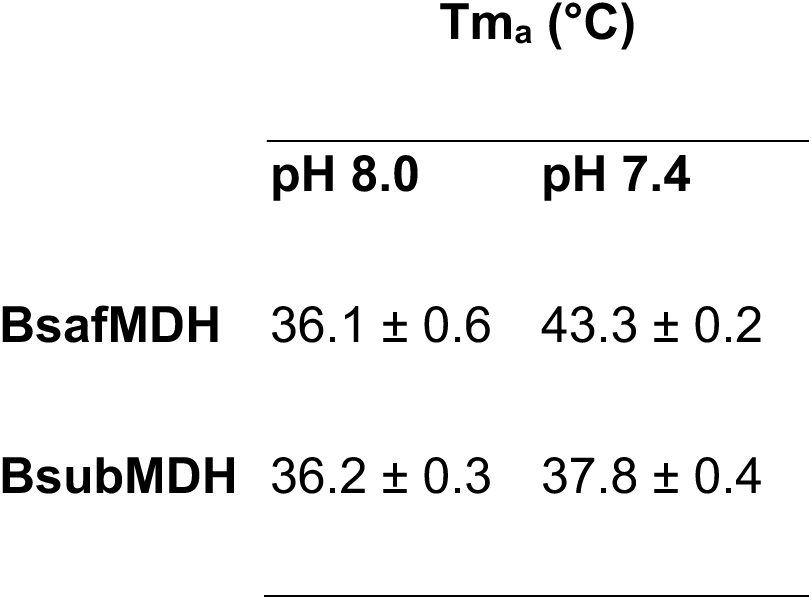
Apparent melting temperatures (Tm_a_s) determined through differential scanning fluorimetry (DSF)

### *B. saf* and *B. sub* MDH are structurally affected by pH

To identify isoform and pH-specific differences in thermal stability and/or conformation, we performed Differential Scanning Fluorimetry (DSF) (also known as fluorescence thermal shift assays). Consistent with the SEC-SAXS results, both enzymes had equivalent apparent melting temperatures (Tm_a_s) at pH 8.0 (36.1 ± 0.6°C for BsafMDH, 36.2 ± 0.3°C for BsubMDH) (Table 3). Upon lowering the pH to 7.4, however, the Tm_a_ of *B. safensis* MDH rose to 43.3 ± 0.2°C, whereas that of *B. subtilis* MDH was elevated only slightly, to 37.8 ± 0.4°C. While BsafMDH and BsubMDH are >90% identical in sequence, we observe that 8 of the 19 positions of difference are differences between charged and uncharged amino acids (Fig. 1). Visualization of the electrostatic surfaces reveals subtle differences in the charge distributions including a larger negatively-charged surface along the central axis of BsubMDH and stronger C-terminal acidic patch in BsafMDH (Fig. 4). These changes in charge do not appear to affect the overall activity of the proteins but do affect their pH stability. These differences in stability evident under laboratory conditions may allow each protein to function optimally in the cellular environment of its respective host.

**Fig. 4.**
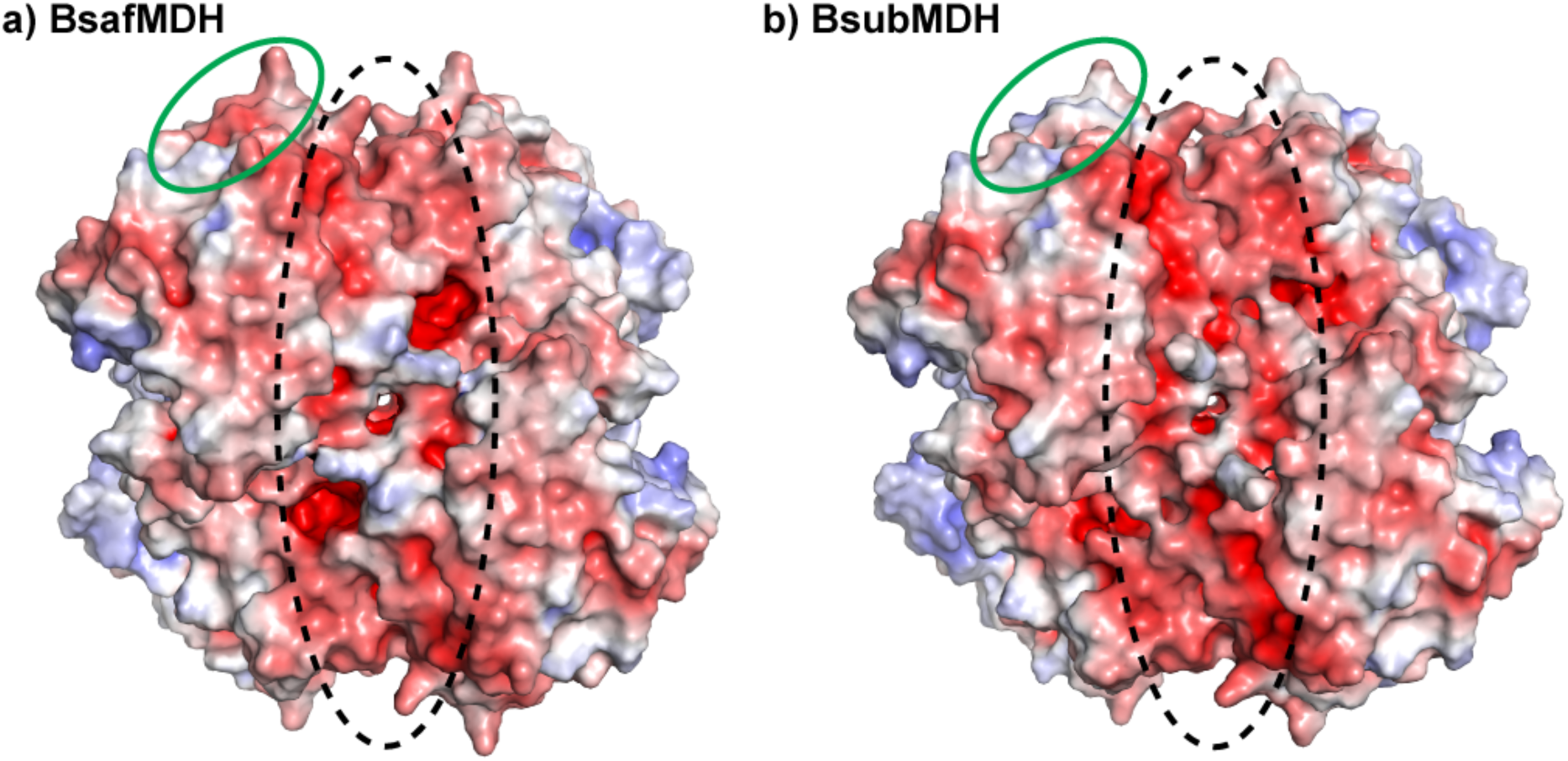
*B. safensis* MDH and *B. subtilis* MDH have different charge distributions. Compared to BsafMDH **(a)**, BsubMDH **(b)** has a more negatively charged surface exposed throughout the central axis between dimers (indicated via black dashed region). BsafMDH, however, has a more acidic C-terminal region, including an acidic patch indicated in green. Electrostatic surfaces for AlphaFold 3 models (Abramson *et al*., 2024) were generated using the APBS Electrostatics plug-in (Jurrus *et al*., 2018) in PyMOL (DeLano, 2002) and displayed on a scale of -5.000 kT/e (red) to 5.000 kT/e (blue).

## Discussion

Malate dehydrogenase (MDH) proteins–best known for their role in the citric acid cycle–are at the heart of central metabolism in most species. Most research on MDH to date has focused on dimeric MDH1 and MDH2 forms of the protein (such as are found in humans), rather than the tetrameric lactate dehydrogenase (LDH)-like form, which contains a dimer of dimers and is found in many species of bacteria (including *Bacillus* species), other prokaryotes, and archaea. This is despite the MDH3/LDH family being more abundant than MDH1 and MDH2 families (Brochier-Armanet and Madern, 2021b) and having uses in both biotechnology and clinical diagnostics (Shimozawa and Nishiya, 2019), in addition to their natural roles. The limited published research on LDH-like MDH proteins is largely focused on MDHs from extremophiles (Richard *et al*., 2000; Lee *et al*., 2001). There remain gaps in our knowledge of the biophysical and enzymatic properties of LDH-like MDHs from more common, and societally relevant, bacteria including *Bacillus subtilis* and *Bacillus safensis*. Here, we show that the MDH proteins from these bacteria are almost identical in their amino acid sequences (Fig. 1), yet they possess subtle differences in biochemical properties. This makes the pair ideal for teasing apart the inner workings of LDH-like MDHs, which can provide much needed information about not only *Bacillus* MDH proteins, but also those of the entire understudied class of LDH-like MDHs.

We found similar enzymatic activity of these enzymes compared to one another in both the reductive direction (oxaloacetate to malate) and the oxidative direction (malate to oxaloacetate) (Fig. 3 and Table 2). Consistent with findings from MDH isozymes from eukaryotic and bacterial MDH throughout the three classes (de Lorenzo *et al*., 2024a), both enzymes showed lower activity in the oxidative direction, with both elevated K_M_ and reduced *k_cat_* values with malate as the substrate. Of note, however, K_M_ values for malate (61 ± 7 µM for BsafMDH and 69 ± 10 µM for BsubMDH), although 5-fold higher than those for OAA, were less than those reported for many MDH isoforms (de Lorenzo *et al*., 2024a). A number of other bacterial MDH have also been found to have comparatively low K_M_ values for malate (Takahashi-Íñiguez *et al*., 2016b). It is possible that this modest K_M_ might provide the versatility required for MDH in prokaryotic cells with single MDH isoforms. However, some prokaryotic MDH show high K_M_ values towards malate (Takahashi-Íñiguez *et al*., 2016b; de Lorenzo *et al*., 2024b).

Despite them having similar shapes and enzymatic activity at pH 8.0, we saw differences between *B. saf* and *B. sub* MDH’s thermal stability at pH 7.4 (Table 3). Although both enzymes had a higher Tm_a_ at the lower pH, suggesting greater thermal stability, the rise in Tm_a_ was larger for BsafMDH than BsubMDH (Tm_a_s of 43.3 ± 0.2°C for BsafMDH and 37.8 ± 0.4°C for BsubMDH at pH 7.4 versus 36.1 ± 0.6°C for BsafMDH and 36.2 ± 0.3°C for BsubMDH at pH 8.0) (Table 3). Decreased pH was also accompanied by increases in both K_M_ (25 ± 3 µM and 11 ± 2 µM for BsafMDH and BsubMDH, respectively, at pH 7.4 compared to 14 ± 3 µM and 8 ± 2 µM at pH 8 .0) and *k_cat_* (180 ± 10 s^-1^ and 109 ± 9 s^-1^ for BsafMDH and BsubMDH, respectively, at pH 7.4 compared to 120 ± 10 s^-1^ and 92 ± 6 s^-1^ at pH 8.0) (Table 2). The differences in activity at pH 7.4 compared to pH 8.0 were greater for BsafMDH than for BsubMDH, but led to similar specificity constants (*k_cat_/*K_M_ = (7 ± 1) x 10^6^ M^-1^ s^-1^ for BsafMDH and (1.0 ± 0.2) x 10^7^ M^-1^ s^-1^ for BsubMDH) (Table 2). These changes are the subject of ongoing investigation in our lab and may be related to differences in charge distribution between the two enzymes (Fig. 4).

Also consistent with what has been reported for a variety of MDH homologs (Takahashi-Íñiguez *et al*., 2016b; Martinez-Vaz *et al*., 2024), we observed substrate inhibition by oxaloacetate (Fig. 3a, 3b, and Table 2). This inhibition was more pronounced at pH 7.4 than pH 8.0, with K_i_ values of 110 ± 10 µM for BsafMDH and 150 ± 30 µM for BsubMDH at pH 7.4 compared to 500 ± 200 µM for BsafMDH and 600 ± 200 µM for BsubMDH at pH 8.0 (Fig. 3a,b and Table 2). Notably, we found this inhibition at lower concentrations of OAA than have been reported for most MDH, suggesting that the *B. safensis* and *B. subtilis* MDH are especially sensitive to substrate inhibition. For example, porcine mitochondrial and cytoplasmic MDH isoforms have K_i_’s towards OAA of 2.0 mM and 4.5 mM, respectively, at pH 7.4 (Bernstein *et al*., 1978). As the authors point out, however, the physiological relevance of this is uncertain, given that levels of OAA in mitochondria are tightly controlled at low levels. Keeping OAA levels low is crucial for maintaining thermodynamic drive for the citric acid cycle to continue in the “forward” (oxidative) direction given the unfavorable ΔG° of the malate → oxaloacetate reaction (Guynn, Gelberg and Veech, 1973). Unlike eukaryotic cells, prokaryotes lack distinct membrane-bound compartments. This may make sensitive regulation all the more important.

In sum, this work provides practical insight into the inner workings of vital understudied enzymes as well as a glimpse into the functional relevance of evolutionary differentiation of MDH proteins into multiple classes. This information could help: 1) answer fundamental basic research questions about protein sequence-structure-function relationships, providing training data for predictive modelling software and 2) enable engineering and efficient use of these enzymes and the bacteria containing them for biotechnological applications including the production of industrial chemicals, medical diagnostics tests, and heavy metal bioremediation.

## Methods

### Recombinant MDH expression

The sequence of *Bacillus safensis* malate dehydrogenase (MDH) was retrieved from UniProt (accession #A0A0M2EAA6), codon-optimized for expression in *E. coli* cells, and cloned into a pET28 vector by Twist Biosciences to add an N-terminal His-tag. A pWH844 plasmid containing the sequence of *Bacillus subtilis* MDH (Uniprot accession #P49814) with an N-terminal His tag (pGP385 (Bartholomae *et al*., 2014)) was obtained as a kind gift from Dr. Jörg Stülke. These recombinant plasmids were transformed into BL21(DE3) and overexpressed through autoinduction in growth media consisting of Lysogeny Broth (LB) supplemented with 0.6% (v/v) glycerol, 0.05% (w/v) glucose, and 0.2% (w/v) lactose. Autoinduced cultures were grown overnight at 30°C with shaking then harvested and resuspended in 50 mM Tris-Cl pH 8.0, 1 mM imidazole, 100 mM NaCl, 0.1 mM EDTA. Resuspended cultures were flash frozen in liquid nitrogen and stored at -80°C prior to subsequent purification.

### Recombinant MDH purification

Thawed resuspended cultures were lysed enzymatically (1 mg/mL lysozyme) and mechanically (via ultrasonication), with the inclusion of Benzonase Nuclease (25 U/µL) and cOmplete EDTA-free Protease Inhibitor Cocktail). Lysates were ultracentrifuged at 17,000 x g for 30 minutes, and the supernatant was subjected to Ni-affinity chromatography. Columns were washed with 50 mM Tris-Cl (pH 8.0), 300 mM NaCl, 10 mM Imidazole, 0.1 mM EDTA, 1 mM phenylmethylsulfonyl fluoride (PMSF). Then, protein was eluted through a gradient to 50 mM Tris-Cl (pH 8.0), 50 mM NaCl, 300 mM imidazole, 0.1 mM EDTA, 1 mM PMSF. Samples containing MDH were pooled and dialyzed overnight against 10 mM Na phosphate buffer, pH 8.0; 50 mM NaCl, 1mM β-mercaptoethanol, 10% (v/v) glycerol. Final concentrations were determined using a BCA assay (Thermo Scientific cat. #23250).

### MDH activity assays

Spectroscopic MDH activity assays were conducted in 1 mL reaction volumes in 50 mM sodium phosphate (NaPi), pH 8 or pH 7.4 with ranges of concentrations of NADH, OAA, and malate. Kinetic parameters for oxaloacetate (OAA) were measured using OAA concentrations from 3.125 µM–200 µM and a fixed concentration of NADH (100 µM). Kinetic parameters for NADH were measured using NADH concentrations from 3.125 µM–200 µM and a fixed concentration of OAA (100 µM). Kinetic parameters for malate were measured using malate concentrations from 7.862 µM–1 mM and a fixed concentration of NAD^+^ (5 mM). Final enzyme concentrations in the assay mixture were BsafMDH: 8.43 nM and BsubMDH: 7.43 nM for assays run in the reductive direction. For assays run in the oxidative direction, final enzyme concentrations were BsafMDH: 12.6 nM and BsubMDH: 7.43 nM.

Reactions were started by the addition of freshly prepared OAA for assays run in the reductive direction and by the addition of MDH for assays run in the oxidative direction. Reaction progress was assessed by continuous monitoring of 340 nm wavelength absorbance, corresponding to NADH concentration, in a diode array UV-Vis spectrophotometer. Each reaction was measured for one minute and the linear portion of each curve was used to determine V_o_ for that concentration of substrate. For NADH and malate assays, data were fitted in R using minpack.lm (Elzhov *et al*., 2023) and nlsLM with the equation rate = (*k_cat_* x [OAA])/(K_M_ + [OAA]) to fit for *k_cat_* and K_M_. For OAA assays, data were fitted in R using minpack.lm (Elzhov *et al*., 2023) and nlsLM with the equation rate = (*k_cat_* x [OAA])/(K_M_ + [OAA] + [OAA]^2^/K_i_) to fit for *k_cat_*, K_M_, and K_i_.

### Differential Scanning Fluorimetry

Thermal stability of BsafMDH and BsubMDH at pH 8.0 and pH 7.4 was determined through differential scanning fluorimetry (DSF) in a BioRAD CFX Duet Real-Time PCR System. 4 µM MDH and 20x SYPRO Orange in 50 mM NaPi (pH 7.4 or 8) were exposed to a temperature gradient from 10°C to 95°C at an interval of 0.5°C per minute with fluorescence scans every 30 seconds. Apparent melting temperatures (Tm_a_s) were determined via curve fitting in DSFWorld (Wu, Gale-Day and Gestwicki, 2024).

### Small-angle X-ray Scattering

Purified protein samples of BsafMDH and BsubMDH were shipped overnight on dry ice and analyzed using SEC-SAXS at the SIBYLS beamline at the Advanced Light Source, Lawrence Berkeley National Laboratory, Berkeley, CA (Classen *et al*., 2013; Rosenberg, Hura and Hammel, 2022). Samples were separated on a Shodex KW-803 column at a flow rate of 0.5 mL/min at 10 °C, and the eluate was measured in line with UV/vis absorbance at 280 nm, multi-angle X-ray scattering (MALS), and SAXS. The incident light wavelength was 1.127 Å at a sample-to-detector distance of 2.1 m. This setup results in scattering vectors, q, ranging from 0.0114 to 0.4 Å^−1^, where the scattering vector is defined as q = 4πsin(θ)/λ, with θ being the measured scattering angle. Radially averaged SAXS data files were processed and analyzed in RAW (Hopkins, 2024). The radius of gyration (Rg) was calculated for each of the subtracted frames using the Guinier approximation: I(q) = I(0) exp(−q2Rg2/3) with the limits qRg < 1.3. The elution peak was compared to the integral of the ratios to background and Rg relative to the recorded frame using the RAW program. Uniform Rg values across an elution peak represent a homogeneous sample. The final merged SAXS profiles, derived by integrating multiple frames at the elution peak, were used for further analysis. We calculated the Guinier plot to provide information on the aggregation state and the pair distribution function [P(r)] to calculate the maximal inter-particle dimension. Models were fitted to the SAXS data using FOXS (Schneidman-Duhovny *et al*., 2013, 2016). All SAXS data have been submitted to the SASDB and the accession numbers will be added prior to publication.

## Supporting information

Supplemental Figure 1 and BsafMDH DNA sequence

## Supplementary Materials

Supplementary Materials contains Figure S1, showing purity of recombinant BsubMDH and BsafMDH enzymes as determined by SDS-PAGE and SEC, and the DNA sequence of the BsafMDH plasmid insert that was cloned into a pET28 vector for recombinant expression.

## Data availability

All SAXS data have been submitted to the SASDB and the accession numbers will be added prior to publication.

## Funding

BB was supported through internal awards from Loyola Marymount University and Saint Mary’s College of California as well as a cohort fellowship from the Malate Dehydrogenase CUREs Community (MCC), with funding through NSF 2119918. CEB and JT were supported by infrastructure provided by NSF MCB RUI-2322867. The SAXS work was conducted in part at the Advanced Light Source (ALS), a national user facility operated by Lawrence Berkeley National Laboratory on behalf of the Department of Energy, Office of Basic Energy Sciences, through the Integrated Diffraction Analysis Technologies (IDAT) program, supported by DOE Office of Biological and Environmental Research. Additional support came from the National Institutes of Health project ALS-ENABLE (P30 GM124169) and a High-End Instrumentation Grant S10OD018483.

## Conflicts of interest

The authors report no conflicts of interest.

## Notes

### Competing Interest Statement

The authors have declared no competing interest.

