## Supplemental Figure 1 and BsafMDH DNA sequence for "Biophysical and enzymatic comparison of Bacillus safensis and Bacillus subtilis malate dehydrogenase (MDH) enzymes"

**Supplementary Materials**

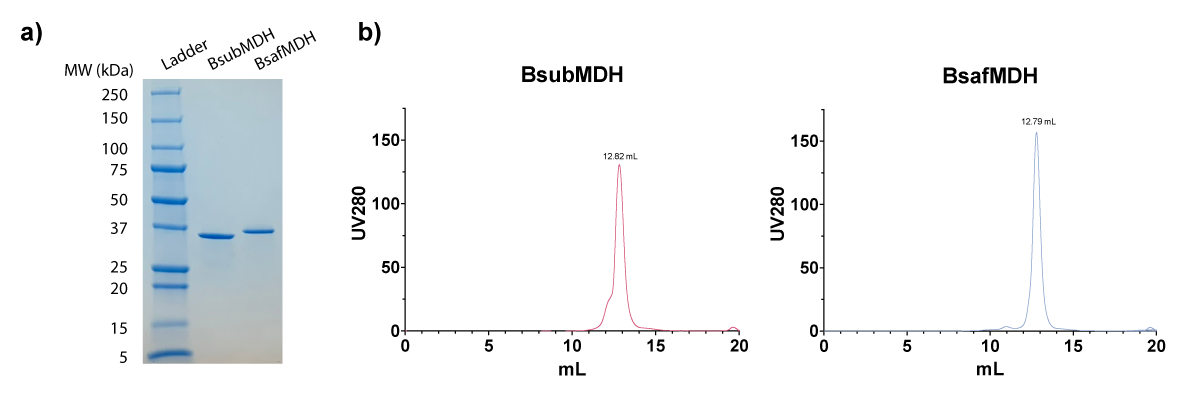

**Figure S1** Purity of recombinant *Bacillus safensis* and *Bacillus subtilis* malate dehydrogenase (MDH) proteins is shown via **a)** SDS-PAGE and **b)** Size Exclusion Chromatography (Superdex 200 Increase 10/300 GL column, Cytiva, Marlborough, MA).

**DNA sequence of BsafMDH clone insert**

The following sequence was cloned into a pET28 vector by Twist Bioscience (Sout San Francisco, CA) with an N-terminal 6XHis tag. It corresponds to the codon-optimized (for *E. coli* expression) sequence of *Bacillus safensis* malate dehydrogenase: Uniprot accession number: A0A0M2EAA6_BACIA.

ATGGCTAATAAAAGGAAGAAAGTATCAGTTATCGGCGCGGGTTTCACCGGTGCGACCACGGCGTTTCTGACCGCGCAAAAAGAACTTGCCGATGTCGTGTTGGTCGACATCCCGCAGCTGGAGAACCCGACCAAAGGTAAAGCGCTGGACATGCTGGAGGCGTCCCCGGTCCAGGGTTTTGACGCTAATATTACCGGCACCAGCAATTACGAAGATACCGCGGGTTCAGATGTGGTTGTGATCACGGCTGGCATCGCCCGTAAACCGGGTATGTCTCGTGATGACCTGGTAAGCACCAACGAAAAAATCATGCGTTCCGTGACGAGAGAAATCGTTAAGTACTCTCCGGAAGCGATCATCGTGGTGCTGACGAACCCGGTTGATGCAATGACCTACGCTGTGTACAAAGAAAGCGGCCTGCCGAAAGAGAAGGTGATTGGTCAGAGCGGTATCCTGGACACCGCGCGTTTCCGCACCTTTGTTGCCCAAGAATTGAATCTGAGCGTCAAGGACGTTACCGGCTTCGTTCTCGGAGGCCACGGCGATGATATGGTTCCGCTCGTTCGTTATAGCTATGCTGGCGGTATTCCTCTGGAGACACTGATTCCGAAGGACCGTATTGATGCGATTGTTGAGCGTACTCGCAAGGGTGGCGGTGAAATTGTTAATTTGCTGGGCAACGGCAGCGCATACTACGCACCGGCAGCAAGCCTGGTTGAGATGGTGGAAGCGATTCTTAAGGACCAGCGTCGTGTTATCCCAACGATCGCGTACTTGGAGGGCGAGTATGGTTATGAGGGTATTTATCTGGGCGTTCCGACTATCGTGGGTGGAAACGGTTTGGAGCAGATTATCGAATTAGAGTTGACCGAAGAAGAGCGCTCGCAACTGGACCGCAGCGTGGAGTCTGTTAAAAACGTGATGAAGGTGCTGTCCTAA
